# Microbial strategies of environmental adaptation revealed by trait-environmental relationships

**DOI:** 10.1101/2024.09.17.613589

**Authors:** Minglei Ren, Ang Hu, Zhonghua Zhao, Xiaolong Yao, Ismael Aaron Kimirei, Lu Zhang, Jianjun Wang

## Abstract

Microbial trait variation along environment gradients is crucial to understanding their ecological adaptation mechanisms. With the increasing availability of microbial genomes, making full use of the genome-based traits to decipher their adaptation strategies becomes promising and urgent. Here, we examined microbial communities in water and sediments of 20 East African lakes with pH values ranging from 7.2 to 10.1 through taxonomic-profiling and genome-centric metagenomics. We identified functional traits important for microbial adaptation to the stresses of alkalinity and salinity based on the significant trait-environment relationships (TERs), including those involved in cytoplasmic pH homeostasis, compatible solute accumulation, cell envelope modification and energy requisition. By integrating these significant-TER traits, we further developed an environmental adaptation index to quantify the species-level adaptive capacity for environmental stresses, such as high pH environments. The adaptation index of pH showed consistently significant positive relationships with species pH optima across regional and global genomic datasets from freshwater, marine and soda lake ecosystems. The generality of the index for quantifying environmental adaptation was demonstrated by showing significant relationships with the species niche optima for the gradients of soil temperature and seawater salinity. These results highlight the importance of trait-environment relationships in facilitating the inference of microbial genomic-based adaptation mechanisms, and expand our understanding of ecological adaptative strategies along environmental gradients.

## Introduction

Microbes inhabit nearly every habitat on Earth and play an important role in biogeochemical cycling as a crucial component of ecosystems (Gilbert et al. 2014). Biogeography and metabolic capabilities of microbial communities have been shown to be driven by multiple environmental gradients occurring naturally in ecosystems, e.g., salinity, temperature, and pH. For instance, the decreasing microbial diversity with salinity is consistently observed across soil (Hollister et al. 2010, Rath et al. 2019) and aquatic ecosystems (Campbell and Kirchman 2013, Mo et al. 2021, Tee et al. 2021). However, our understanding of functional mechanisms underlying the diversity patterns along environmental gradients lags behind. Generally, environmental condition affects the growth of microbes or their competition, and selects directly or indirectly against microbial lineages with variable adaptative capability to environmental changes (Cadotte and Tucker 2017). Resolving the succession in species adaptation potential along environmental gradients will greatly improve our understanding of biogeography and habitat expansion for microbes.

The trait-based concept, which focuses on the fitness-influencing characteristics of individuals, is widely applied to identify the underlying mechanism driving community assembly and dynamics. (Martiny et al. 2015, Yang 2021, Wang et al. 2022). For instance, the traits inferred from microbial genomes or proteomes reflect a cascade of cellular and molecular regulation, and potentially predict species ecological performance (Hu et al. 2021, Ren and Wang 2021, Westoby et al. 2021) and their adaptive strategies to aridity (Li et al. 2022). Microbial traits are further used to explore their life history strategies, including the copiotroph-oligotroph (Fierer et al. 2007), the competitor-stress tolerator-ruderal (Ho et al. 2013, Krause et al. 2014) and the growth yield-resource acquisition-stress tolerance strategies (Malik et al. 2020).

How to identify the traits that are linked to microbial life strategies or their adaptive mechanisms remain a challenge. The trait-environment relationships (TERs), that is how microbial traits relate to environments, are generally used to inform the traits linked to environmental changes (McGill et al. 2006, Funk et al. 2017), enhancing our understanding of functional dynamics of microbial community along environmental gradients. Such traits identified through the similar approaches could be utilized to inform environmental adaptation for microbes. For instance, the species pH preference could be predicted through a model of the presence or absence patterns of the pH-associated traits, which are identified from representative genomes linked to conserved marker genes (Ramoneda et al. 2023). Even more complex microbial functions, including salinity preference and methanogenesis, are commonly summarized by a combination of multiple traits, rather than captured by a single trait (Martiny et al. 2015). Therefore, how to define quantitatively microbial adaptation ability considering the joint effects of multiple functional traits is pivotal for predicting the responses of microbial communities to these environmental changes and their implications at the global scale.

In this study, we examined the functional trait patterns of microbial communities in water and sediments of African lakes along a pH gradient, and developed a trait-based index to quantify species’ capacity for environmental adaptation through a framework of three steps (Figure 1). Firstly, we determined the TER relationships between the community-weighted means of functional traits that are represented by KEGG orthologs (KOs) and lake pH by the effect size metrics, such as Spearman correlation coefficient. We secondly identified the functional traits important for microbial adaptation based on the significant TERs, and finally presented a novel index of a species to quantify how to adapt to an environmental stressor by calculating the mean value of the TERs across species traits weighted by the copy number of KOs. Our novel approaches and main findings in the study suggest that the utilization of functional traits inferred from microbial genomes not only expectedly move beyond descriptive patterns in community composition, but also improve our understanding of genetic mechanisms driving biogeographic distribution and their adaptation to global environemtnal change.

**Figure 1.**
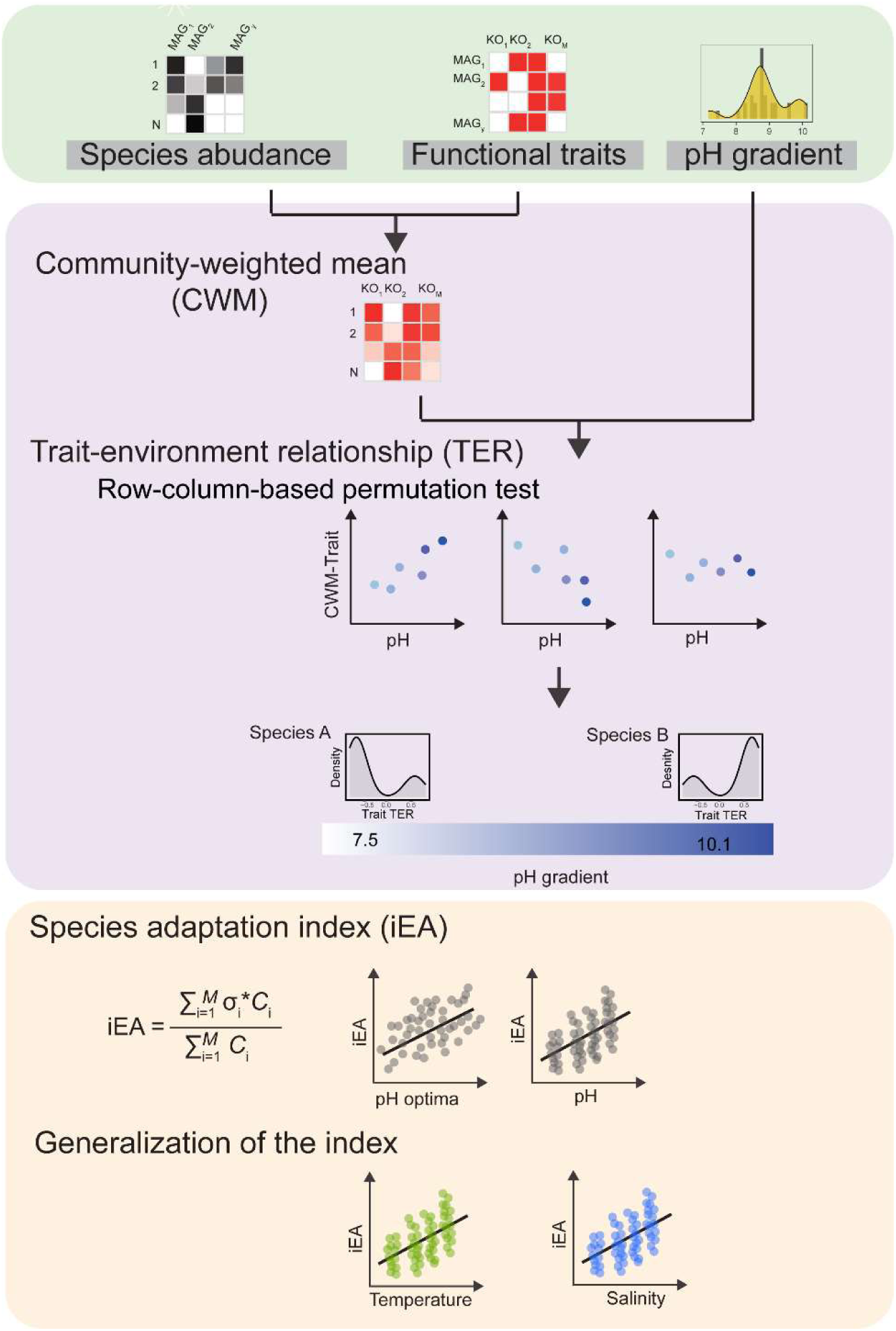
The overview of the trait-based framework in the study. The overflow of the framework consists of three modules, highlighted with rectangular boxes in distinct colors. At the top module, two data tables were prepared through taxonomic profiling and metagenomic-centric analyses, including the species abundance table and functional traits tables. At the middle module, we identified functional traits important for microbial adaptation through trait and environment relationship (TER), and then presented an index of environmental adaptation to quantify species adaptive capacity to the environmental stressor at the bottom. TER was determined by the effect size metrics to represent the strength of the relationship between microbial trait and environmental stresses. The details about each step see the Materials and Methods.

## Materials and Methods

### Field sampling

We collected a total of 39 samples, including 19 water and 20 sediment samples, from 20 lakes and reservoirs in East Africa during February 2020 (Appendix S1: Figure S1). These lakes have a wide range of salinities (56∼85,318 ppm) and pH (7.2∼10.1). More detailed description about the sampling sites is found in previous literature (Yao et al. 2022, Zhao et al. 2022). Total DNA of samples was extracted using the PowerSoil DNeasy Kit (QIAGEN, Germany) under sterile conditions, and then subjected to metagenomic sequencing according to the manufacturer’s protocol (Magigene, China). The information about sample collection, DNA extraction and sequencing is detailed in the Supplementary Methods and Materials. The physicochemical parameters are shown in the FigShare Table S1, with the DOI number 10.6084/m9.figshare.24196512.

### Taxonomic profiling using metagenomic sequencing

Taxonomic profiling of microbial communities across samples was performed based on metagenmic sequencing using a custom pipeline established recently (Ren and Wang 2022). Breifly, the raw metagenomic reads were cleaned, trimmed, and assembled into contigs for all samples. The metagenome-assembled genomes (MAGs) were finally recovered from each of these assemblies. The detailed information about sequence cleaning, assembly and MAG recovery sees the Supplementary Methods and Materials. The statistics regarding clean reads and metagenomic contigs for 39 sample are shown in FigShare Table S2. The information of 1,001 high-quality nonredundant MAGs is shown in FigShare Table S3.

There were 2,677 representative species identified from assembled contigs of all samples based on the conserved ribosomal protein rpS3 using a modified pipeline (Diamond et al. 2019). The species here is defined as the representative of ribosomal protein-coding *rpS3* genes that clustered at the 99% similarity (Sharon et al. 2015). The abundance of these representative species in a sample was calculated as the total bases mapped on the representative sequence divided by the length of the representative sequence and the total number of sequence in the corresponding sample. The taxonomy for these species was determined by sequence alignment against a custom *rpS3* reference database, which was derived from NCBI RefSeq prokaryotic genome database. The taxonomy was further validated and carefully corrected according to phylogenetic tree of rpS3 genes identified above. The detailed information about species identification, abundance estimation and taxonomic assignment sees the Supplementary Methods and Materials.

The genomes for representative species were retrieved to obtain functional traits for microbes through the *rpS3*-containing sequence shared between the reconstructed MAGs and the *rpS3*-represented species, resulting in 647 representative genomes. The functional traits were represented as orthologous genes from the most commonly used annotation databases, such as KEGG orthologs (KOs). The KOs annotation links the orthologs to molecular-level functions and higher-level metabolic processes that provide a high resolution for the biochemical role of traits (Kanehisa et al. 2016). The protein-coding gene sequence of each representative genome was predicted by Prodigal v2.6.3 (Hyatt et al. 2010), then annotated against the eggNOG database v5.0 (Huerta-Cepas et al. 2018) using eggNOG mapper v2.1.6 with DIAMOND as the seed ortholog search engine (Cantalapiedra et al. 2021). The assignments of KEGG orthologs (KOs) for each species were retrieved from annotation results of eggNOG mapper using custom Perl scripts.

### Development of the functional trait-based framework

A trait-based framework was developed to qualify the species capacity to adapt to environmental stresses by considering its key functional traits and their relationships with environments. There were three primary steps in the framework, which were illustrated based on the microbial communities in African lakes along a pH gradient as below. Three dataset tables were used in the following analyses: (1) the species abundance table with 2,677 species from 39 samples; (2) a list of environmental factors for 39 samples; (3) the functional traits represented by KOs for 647 representative species with genome available. All statistical analyses were performed in the R language v4.1.1.

Firstly, the trait and environment relationship (TER) was determined by effect size metrics. Specifically, the effect size could be calculated by the Spearman correlation coefficient rho between a community-weighted mean value (CWM) of trait and pH across samples. The CWM values of each KO were calculated with species abundance across samples and KO copy numbers or genomic traits using the ‘functcomp’ method in the FD package v1.0-12.1 (Laliberté and Legendre 2010). A variety of statistical tests have been developed to assess the trait-environment relationships since last century (Grime 1974), and could be summarized into two categories, namely the species-level and community-level approaches (Kleyer et al. 2012, Lepš and de Bello 2023). The former approach is asking whether the species’ trait relate to their preference along a gradient, and the latter is about how trait distribution within community changes along a gradient. There are two important issues to be usually criticized: trait values of a few dominant species could strongly influence the variation of CWM traits, and the repetitive presence of similar species composition within communities could cause changes of any considered traits to be significant.

Accordingly, the methods with specific null models are developed to correct the potential Type I error rates (Peres-Neto et al. 2017, Zelený 2018, ter Braak 2019). Among these methods, we selected the combination of two independent permutation test strategies. One is the row-based permutation randomizing trait values in the community-level approach and testing the species attributes-species composition link. The other is the column-based permutation randomizing environmental variables for the species-level method and testing the sample attributes-species composition link. The conservative *P*-values of both two tests were chosen as a result. The combined permutation strategy has been demonstrated to control the type I error across aquatic and terrestrial environments (Peres-Neto et al. 2017, Zelený 2018). Due to the word limitation, the detailed discussion and additional tests regarding the selection of the permutation strategy see the Supplementary Method. The strategies implemented by the function ‘test_cwm’ in the ‘weimea’ package v0.1.10 was used to analyze the trait and environment relationship.

Secondly, potential adaptive mechanisms were inferred by the genomic and functional traits significantly correlated with pH. The functional traits important for microbial adaptation were identified based on the significant TERs. The important roles of selected traits in mediating species adaptation to environment stresses was confirmed by the consistency of these functions with expert opinions or current knowledge in literature (FigShare Tables S4 and S5). Further inspection of the other significant functional traits with curated annotation was applied to provide novel insights into putatively adaptive mechanisms but rarely reported in previous literature.

Finally, we presented an index of environmental adaptation (iEA) to quantify the species’ adaptive capacity for environmental stresses by considering its key traits and their relationships with environments (TERs). The iEA was calculated by the formula 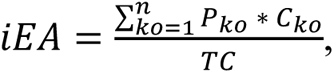, where *P_ko_* is the coefficient of Spearman rank correlation between pH and a functional trait represented by KO; *C_ko_*, copy number of the KO, and *TC*, the total count of KO copy number in the species. The extent of the iEA to the species optimum niche was accessed by the relationship between the index of a species and its pH optima through a linear regression model. The pH optima of an individual could reflect the optimum niche where the species is expected to be most abundant, and was calculated based on its abundance across samples and lake pH of samples where the species was identified. The formula was shown as 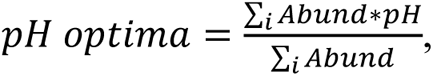, where ‘*Abund*’ is the abundance of the species in sample *i* and ‘*pH*’ is the pH of sample *i*.

### Validation and generalization of the index

The utility of the adaptation index for microbial species was validated using three independent genome datasets with explicit pH gradients. (i) The first one is a set of 345 MAGs, which were reconstructed in the study, but not included in any of data analyses above. (ii) The second dataset consists of 530, 464 and 606 species-level MAGs from freshwater, marine and alkaline habitats, respectively. (iii) The third dataset contains 830 high-quality species-level MAGs from freshwater and soda lakes with pH values available. The accession number and relevant environmental information of the last two datasets were shown in the FigShare Tables S6 and S7. The detailed information about these three genomic datasets sees the Supplementary Methods and Materials.

The generality of the framework for other environmental gradients was examined by two sets of metagenomic datasets along salinity and temperature gradients. The first dataset contains 12 soil samples collected from a slow-burning coal-seam fire around Pennsylvania (USA), with soil temperatures ranging from 13.3 to 54.2 ℃ (Sorensen et al. 2019). The second comprises 10 surface marine water samples followed by a 0.2 µm filtration, from the Baltic Sea with salinities ranging from 2,440 to 28,100 ppm (Alneberg et al. 2018). The procedures of performing sequence analysis and trait-based framework were the same as the cases of the African microbial community along a pH gradient above. The accession number and relevant metadata for the samples in both gradient datasets are shown in the FigShare Table S8.

## Results

### Multilevel adaptive strategies revealed by the trait-environmental relationships

Before investigating how functional traits relate to pH gradient, we identified pH to be an important environmental driver of the lake microbiomes based on the following analyses. (i) The Mantel test analysis indicated that microbial communities were significantly correlated with the difference in lake pH with the highest Mantel r value of 0.35 and 0.34 for water and sediment communities, respectively (*P* < 0.0001, Mantel test, Appendix S1: Figure S2a). (ii) The random forest model result revealed that pH was most important among measured environment variables for the Shannon diversity of water communities, and water temperature for sediment communities (Appendix S1: Figure 2b).

**Figure 2.**
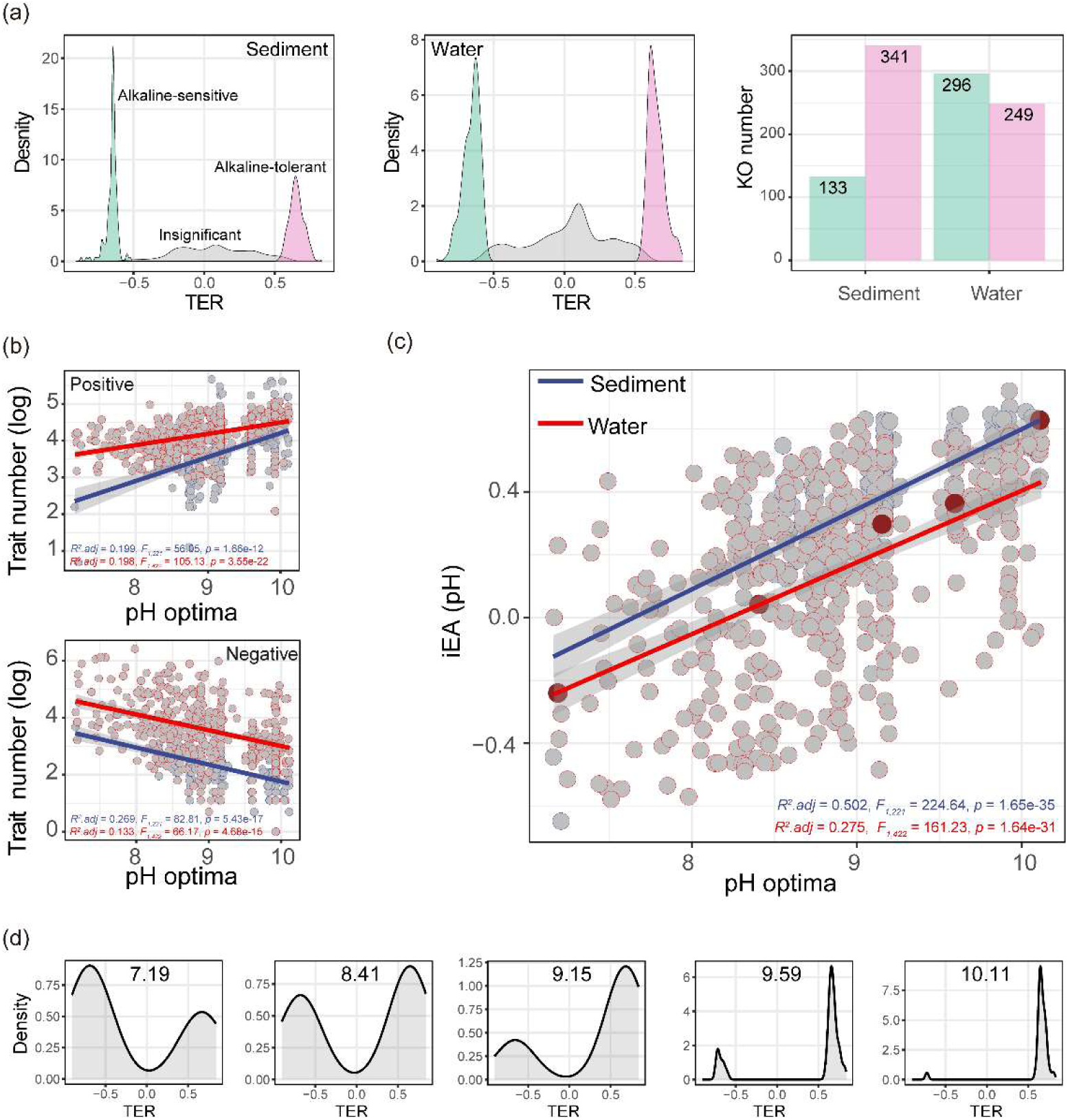
The index of environmental adaptation and their associations with species niche optimum. (a) The distribution of TER for all functional traits across distinct habitats. Functional traits were classified into three groups based on the significance of TERs as shown in the plot. TERs were tested by the Spearman correlation test followed by the row-and column-based permutation test. (b) The number of alkaline-sensitive traits occurring in species decreased with their pH optima, and the number of alkaline -tolerant traits increased with their pH optima. (c) The relationships between the species iEA index and pH optima across water and sediments were fitted using a linear regression model. (d) Five species from the phylum Chloroflexi were selected to illustrate the reorganization of alkaline-tolerant or alkaline-sensitive traits in microbial species for their adaptation to high pH stress.

We identified the functional traits important for microbial adaptation based on the significant relationships between the community-weighted means of traits and lake pH, that is, the trait-environment relationships (TERs, Table 1, see the details in Materials and Methods). Briefly, the TERs were examined with Spearman rank correlation, and the significance was tested with a row-and column-based permutation test (Zelený 2018) to control inflated Type I error in the statistical analyses of TERs (Peres-Neto et al. 2017). We found that there were 952 significantly pH-associated functional traits represented by KEGG orthologs (KOs), including 478 KOs for planktonic microbes, 407 KOs for benthic microbes, and 67 KOs shared by both communities (Figure 2a, FigShare Table S4). The number of these KOs only accounted for 14.4% of all 6,600 KOs identified from all 647 representative genomes from African lakes, while the other 5,648 KOs were nonsignificantly correlated with pH. When inspecting the distribution of these genes across individuals, the number of these pH-related genes ranges from 10 to 391 for each species, with the median value of 137 that accounts for 5.32% of all protein-coding genes.

**Table 1.** Glossary of the relevant terms referred in the study.

| Terms | Explanation |
| --- | --- |
| Environmental adaptation | The adjustment of an organism’s features in order to enhance its survival in specific environments, which have been shaped by evolutionary forces. |
| Functional traits | The characteristics that affect the fitness of a microorganism under certain environmental conditions. They could be defined as orthologous gene families encoded by microbial genomes, such as KEGG orthologs (KOs). |
| Community-weighted mean (CWM) | A community-level aggregated measure of traits, reflecting mean features of microbial communities through combining the information of species traits and their relative abundance. |
| Trait-environment relationship (TER) | TER describes the relationship between the community-weighted mean values of traits and environment conditions. For a functional trait, the strength and sign of TER are determined by effect size metrics, such as Spearman rank correlation coefficient between a community-weighted mean of the trait and a major environmental factor. The strength of TER is represented by the absolute value of correlation coefficient, whereas the sign of TER informs whether the trait is for or against environmental stress. |
| Index of environmental adaptation (iEA) | The iEA is to quantify a microbial species capacity to adapt to an environmental stressor by considering its key traits and their relationships with the environment (that is, TER). The iEA is calculated as the mean value of the TERs across species traits weighted by the copy number of KOs. |

Based on the significant TERs, we further identified two fundamental properties for each functional trait: strength and sign. Specifically, the strength of TER is represented by the absolute values of correlation coefficients, and indicates how much a functional trait could contribute to the microbial adaptation to pH stress. For instance, an absolute Spearman rho of 1 indicates constant increase or decrease of traits at the community level along a pH gradient, while 0 indicates no variations. The signs of TER inform whether the trait is for or against pH stress, and the positive and negative correlation coefficients indicate the alkaline-tolerant and alkaline-sensitive traits, respectively.

Among the 952 functional traits, there were 545 alkaline-tolerant traits with their TER strength ranging from 0.530 to 0.836, 384 alkaline-sensitive traits with their strength ranging from 0.524 to 0.906, and 23 traits with contrasting TER signs between water and sediments (Figure 2a, FigShare Table S4). Among these functional traits, there were at least 97 KO-represented traits (FigShare Table S5), which are well-known for their involvement in microbial pH adaptation (Sorokin et al. 2014, Banciu and Muntyan 2015). These traits could be grouped into four main strategies, including cytoplasmic pH homeostasis, cell envelope modification, compatible solute accumulation, and energy acquisition. For example, 22 functional traits associated with cytoplasmic pH homeostasis showing significant positive relationship with pH, including the Mrp-type Na^+^:H^+^ antiporter (*mrpABCDEFG*) and the monovalent cation: H^+^ antiporter of the CPA1 family. There were at least 21 well-annotated traits relevant to the biosynthesis and uptake of compatible solutes showing significant positive relationships with pH (*P* < 0.05), including biosynthesis of ectoine (i.e., *asd*, *ectBC*), trehalose (i.e., *otsAB*, *treP*), and betaine (i.e., *betB*, *cmo*), and a variety of transport system such as osmoprotectant ABC transport system (*opuABC*) and the glycine betaine/proline ABC transport system (*proVWX*).

### A TER-based index to quantify how microbes adapt to environmental stress

We further presented an index of environmental adaptation (iEA) to quantify the adaptive capacity of a species to environmental stress by considering its key traits and their relationships with environments (Figure 1, Table 1). Briefly, the iEA is calculated as mean value of the TER across the species’ traits weighted by the copy number of KOs, and these means could be further statistically related to the environments that the species survive such as by Spearman correlation or linear regression. The iEA is an unitless metric regardless of environment factors considered, and has values ranging from -1 to 1. Using lake pH as an example, a species with higher iEA has stronger adaptation capacity for high pH environments whereas lower value of iEA indicates that the species is more sensitive to high pH. Positive and negative iEA values indicate that a microbial species is dominated by alkaline-tolerant and alkaline-sensitive traits, respectively, while an iEA of zero suggests the two kinds of traits equally dominate. More details about the calculation of the iEA were shown in Materials and Methods.

For the microbes in African lakes, the iEAs of pH had considerable variations ranging from -0.648 to 0.721 across habitats and phyla. The microbes had lower levels of iEA in water (0.144 ± 0.301) than sediments (0.375±0.184; Wilcoxon rank sum test, *W* = 69360, *p* = 1.47e-22), although the species iEAs of pH showed similar ranges between two habitats. Moreover, the iEAs of pH showed distinct distributions across phyla in lakes, such as the phylum Bacteroidetes with 0.471 ± 0.071 and Betaproteobacteria with -0.315 ± 0.203 (Appendix S1: Figure S3a). The changes in the species iEAs of pH across habitats and phyla are likely associated with the difference in trait number and TER strength of species, which could reflect their distinct lifestyle across habitats and evolutionary histories among phyla.

Interestingly, we found that the species iEAs of pH were significantly positively correlated with their pH optima across the water and sediments with the *R^2^* of 0.275 and 0.502, respectively (Figure 2c). These correlations confirmed that the iEA could quantitatively represent species adaptive capacities to the studied environmental gradient (i.e., pH). Ecological meanings of the correlations could be explained by a clear reorganization of alkaline-tolerant or alkaline-sensitive traits in microbial species for their adaptation to high pH stress, which are illustrated by the distribution of functional traits (i.e. their sign and strengths) in the five species selected from the phylum Chloroflexi along the pH optima gradient (Figure 2d). For instance, the species with the lowest and highest pH optima of 7.19 and 10.11 had iEA values of -0.240 and 0.628, respectively, and expectedly showed contrasting percentages of alkaline-tolerant traits among all traits that significantly associated with lake pH. The former species contained 36.8% alkaline-tolerant traits, while the latter had nearly tow times higher and reached 97.0%. Consistently, the Chloroflexi species with a lower iEA are mostly found in a neutral pH condition, such as aquifer sediments or deep lakes (Hug et al. 2016, Mehrshad et al. 2018), whereas the species with a higher iEA are reported in alkaline hot springs (Hanada et al. 2002, Meer et al. 2010). We noted that these two species had similar TER strength, that is 0.683 ± 0.075 and 0.674 ± 0.044, respectively. These examples highlight the important roles of the reorganization of distinct traits (i.e. alkaline-tolerant and alkaline-sensitive) and their TER strength for environmental adaptation along gradients.

Further, the species iEAs of pH were also correlated with their pH optima across nine dominant phyla such as Chloroflexi and Alphaproteobacteria. For instance, the bacteria within the phylum Chloroflexi significantly associated with pH optima and showed the highest explained variation with *R^2^_adj_* of 0.783 (*F_1,31_* = 116.38, *p* = 5.08e-12), whereas Alphaproteobacteria had the lowest explained variation with *R^2^* of 0.191 (*F_1,40_* = 10.67, *p* = 2.24e-3, Appendix S1: Figure S3b). These differences in iEA values and their explained variation in pH optima among phyla are likely explained by their preferring adaptation strategies associated with distinct functional traits. These results not only suggest that the species iEA of pH appears to be phylogenetically conserved (Martiny et al. 2015), but also provide additional evidence for the quantification of species environmental adaptation using iEA at the broad level of taxonomic classification.

It should be noted that the number of traits in microbes was also found to be important for their pH optima, but the inclusion of the TER strength in iEA could enhance its predictive power for species niche optima than trait number. We found that the species with more alkaline-tolerant traits exhibited the higher pH optima, whereas the species harboring more alkaline-sensitive traits had relatively lower pH optima (Figure 2b). Specifically, the number of alkaline-tolerant traits that each species carries was positively correlated with their pH optima across water and sediments with *R^2^* of 0.198 and 0.199 respectively (Figure 2b), and the number of alkaline-sensitive traits was negatively correlated with pH optima in water and sediments with *R^2^_adj_* of 0.133 and 0.269, respectively (Figure 2b). However, the trait number showed a relatively lower performance in explaining the variation in pH optima compared with the iEA considering the TER strength, highlighting the important role of TER strengths in predicting the species niche optima.

### Validating the utility and generality of the index of environmental adaptation

To validate the utility of the iEA and demonstrate the independence of iEA on pH, we applied to three genome datasets with explicit pH gradients at both region and global scales (FigShare Tables 6 and S7). The first dataset was the redundant genomes (n = 345) reconstructed from the same metagenomes of the African lakes as the representative genomes (647), but not used anywhere, including the KO identification and CWM calculation. When we calculated the iEA for these redundant genomes based on the selected KOs (972) and their relevant TER values, we found that the iEA still correlated with the pH of the samples from which these species were present (Figure 3a). The second dataset consists of the genomes (1600) from global freshwater, marine and soda lakes (Zhao et al. 2020). Based on the same set of selected KOs, we calculated the iEA for these independent genomes, and found that the iEA for the species in soda lakes was significantly higher than other aquatic environments (Figure 3b). The last genome dataset was retrieved from lakes with pH values ranging from 4.1 to 10.3 (Buck et al. 2021). Based on the selected KOs, we found that the iEA for these genomes also correlated with lake pH with *R^2^_adj_* of 0.130 (Figure 3c).

**Figure 3.**
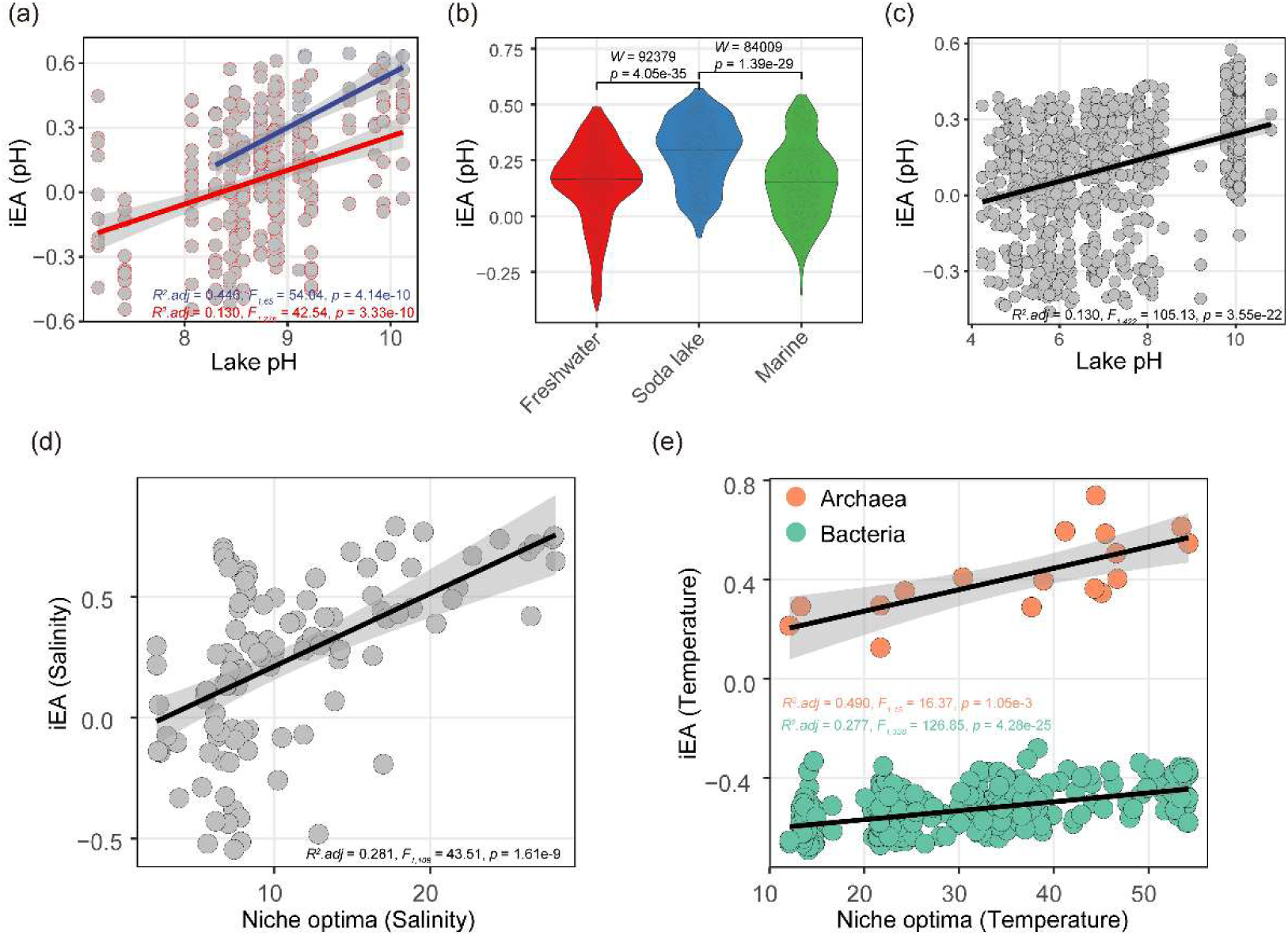
The validation and generalization of strong association between the species iEA and their niche optimum. The relationships between the species iEA index and pH optima or lake pH were validated across three regional (a) or global genomic datasets (b, c). The generality of the species adaptation index across a soil temperature gradient (d) and a marine salinity gradient (e). The iEA of temperature for soil bacteria and archaea was plotted separately against niche optima. The calculation of species iEA in (a, b, c) was the same as that for pH, using the same TER strength and sign derived from the East African lake, whereas the calculation of species iEA of two gradients (d, e) was the same as that for pH in the study. The relationship between adaptation index and niche optimal was fitted using a linear regression model (a, c, d, e) and the significance of difference in the index across habitats was tested by Wilcoxon signed rank test (b). The complete statistical reporting for these tests were shown on each subpanel. Detailed information about three validation genomic datasets of a pH gradient sees the FigShare Tables S6 and S7, and detailed information about two datasets of salinity and temperature gradients sees the FigShare Table S8.

To further support the generality of this index, we extended to another two environmental gradients (FigShare Table S8), namely a salinity gradient ranging from 2.44 to 28.1 ppt in a seawater dataset (Alneberg et al. 2018) and a temperature gradient ranging from 13.3 to 54.2 ℃ of fire-affected soil data (Sorensen et al. 2019). On the one hand, functional traits significantly associated with temperature or salinity were identified using our approach, including the ones known in original studies and the novel ones rarely discussed previously. For instance, the abundance of functional traits involved in sulfur metabolism increased toward hot soils, whereas the abundance of traits involved in two-component regulatory systems decreased, which are consistent with previous findings (Sorensen et al. 2019). The novel traits enriched in the thermophiles in hot soils, including the CRISPR-associated protein that provides adaptive immunity against phages in thermophilic bacteria (Paez-Espino et al. 2015, Munson-McGee et al. 2018) and spore biogenesis and germination, may also play key roles in the resistance of mesophiles and/or thermophiles to thermal stress (Huang and Hull 2017, Ursem et al. 2021). On the other hand, the species iEA of salinity and temperature also showed positive relationships with their corresponding niche optima, with the *R^2^_adj_* of 0.281 for the seawater salinity gradient (Figure 3d), and the *R^2^_adj_* of 0.490 and 0.277 for soil bacteria and archaea along the soil temperature gradient, respectively (Figure 3e). Collectively, these consistent relationships between the indices and niche optima across multiple environmental factors indicate that we could potentially extend the index to quantify species adaptation capacity along any environmental gradients of interest.

## Discussion

Our study presented a novel index to quantify microbial adaptive capacity to environmental stresses by integrating both the strength and direction of the relationships between their functional traits and the driving environment. The trait-based index of environmental adaptation has important implications in predicting taxon-specific biogeographic distribution and their response to global climate change given the high proportions of uncultured species (Steen et al. 2019) and the increasing number of available microbial genomes.

The adaptation index has a robust ability to evaluate species’ adaptive capacity for environment stress as shown by its consistent associations with species niche optima or environments across freshwater, marine and soil ecosystems. The evaluation of species adaptive capacity based on the index could help guide how to optimize media formulations for uncultivated taxa with the increasing expansion of genomic data (Lewis et al. 2021). The index could be used as a quantitative traits of microorganisms to be complementary for their limited phenotypic traits (Westoby et al. 2021, Wang et al. 2022), and be further incorporated as a species key attributes in meta-analyses or modelling across ecosystems (Allison 2012, Barberán et al. 2014).

The adaptation index could be generalized to typical environmental factors across ecosystems, such as pH, temperature and salinity as shown in the study. Since the index is an unitless and standardized metric with its values ranging from -1 and 1, regardless of environment factors, we could easily compare the indices concerning a single environment or even the component of multiple environmental factors (i.e. PCA) within a study. Taking the microbes in the African lake as an example, we calculated the indices of all other environmental factors for the species in the same way as of pH. As expected, these indices showed distinct distributions within a constrained range (Appendix S1: Figure S4a) and significantly associated with the species niche optima of the corresponding factor (Appendix S1: Figure S4b). Interestingly, we found that, compared to other iEA indices, the index of pH had the highest correlation with species pH optima showing a *R^2^_adj_* of 0.332 (Appendix S1: Figure S4c). The results demonstrate the generality of the index for various environmental variables, and also confirm a major role of pH in shaping taxonomic and functional compositions of microbial communities in the African lakes.

We further expect that the establishment of a quantitative environment-associated trait database could be helpful to explore microbial adaptation capacities along environmental gradients across different ecosystems. The functional traits, represented by KEGG orthologs and their relationships with the driving environmental factors could be generally transferred to a wide range of studies to quantify microbial adaptive capacity across spatial scales. This was demonstrated by our three independent validation datasets where the iEA indices of pH were calculated using the TERs derived from the African lakes with a pH gradient. As the increasing number of ecologically relevant traits from microbial genome sequences (Madin et al. 2020, Cébron et al. 2021, Karaoz and Brodie 2022), we expect that the efforts for establishing a database of functional traits with both strength and signs in their relationships with typical environments, such as temperature and nutrients, would facilitate to evaluate the independent or additive responses of the species to global climate changes.

## Acknowledgements

We sincerely thank Huruma Mgana, Mary A. Kishe, Athanas Mbonde, and Omari Wijuru for their assistance in field sampling and laboratory analyses. We would like to thank the Tanzania Commission for Science and Technology (COSTECH) for the research clearance and permit (2019-609-NA-2018-238) to conduct this study and the Tanzania Fisheries Research Institute (TAFIRI) for coordinating the field research.

## Authors’ contributions

The research was conceived by JW. Sample collection was performed by LZ, XY, and JW. Data analyses and drafting of the manuscript were undertaken by MR with the contribution of JW. All authors were involved in the discussion and approved the final version of the manuscript.

