## Supplemental material and figures for "Microbial strategies of environmental adaptation revealed by trait-environmental relationships"

(1) Supplementary materials and methods

(2) Supplementary figures

**Supplementary materials and methods**

**Sample collection, DNA extraction and sequencing**

For each site, surface sediments of 0-5 cm were collected using a bottom sampler and then sealed immediately in the airtight plastic bags. Surface water of 1.5 L was sampled from ~20 cm depth, stored in two sterile polyethylene terephthalate bottles, and filtered through 0.22 µm polycarbonate membrane (Millipore Sigma, Burlington, USA) within 8 hours after collection. Both water and sediment samples were collected near the shores of the lakes, stored in a dark cooler box with ice and then transported to the laboratory at -4 ℃. Sediment samples were freeze-dried at -50 ℃ and finally stored at -20 ℃ in the laboratory.

Total DNA for all samples was extracted using the PowerSoil DNeasy Kit (QIAGEN, Germany) under sterile conditions. Approximately 0.4 g of dried soil was used for DNA extraction in sediment samples, whereas filtering membranes with enriched microorganisms were used for water samples. DNA was subjected to metagenomic sequencing according to the manufacturer’s protocol (Magigene, China). Genomic DNA was first fragmented using ultrasonication, and fragments ranging from 250 bp to 350 bp were targeted. The libraries were prepared using the NEB Next Ultra DNA Library Prep Kit and sequenced on the Illumina NovaSeq6000 platform using 2*150 bp paired-end sequencing strategy. The metagenomic dataset was then subjected to a custom pipeline as described previously (Ren and Wang 2022). Briefly, the quality of raw reads was checked using FastQC v0.11.8 (Andrews 2010), which verified that there were no sequencing adapters in the samples. The raw reads were then trimmed using Trimmomatic v0.39 (Bolger et al. 2014), discarding the reads with an average Phred quality score lower than 25 using a 4-bp-wide sliding window and reads shorter than 50 bp. The clean reads of all samples were prepared for downstream analysis for the different purposes described below (Fig. 1).

**Metagenomic assembly and binning**

*De novo* assembly of metagenomic reads for each sample was performed by MEGAHIT v1.2.8 (Li et al. 2015), a memory-efficient and very fast metagenomic assembler, with the parameter ‘--presets meta-large --min-contig-len 1000’. DAS Tool v1.1.1 (Sieber et al. 2018) was applied to recover metagenome-assembled genomes (MAGs) from each assembly, which determines the optimized MAGs through a strategy of dereplicating, aggregating and scoring the binning results from multiple widely used binning tools. Before recovering microbial genomes, clean reads of each sample were mapped against assembled contigs from the sample using Bowtie2 v2.5.0 with default settings (Langmead and Salzberg 2012), generating an alignment file for the following binning programs. Four metagenomic binning programs were first used to reconstruct preliminary MAGs from the metagenomic assembly, including MaxBin2 v2.2.6 (Wu et al. 2016), MetaBat1 v0.24.1 (Kang et al. 2015), MetaBat2 v2.12.1 (Kang et al. 2019) and CONCOCT v1.1.0 (Alneberg et al. 2014). Then, these binning results were integrated by the DAS Tool, resulting in a final list of dereplicated MAGs. The completeness, contamination and taxonomic assignment for these MAGs were evaluated using CheckM v1.0.13 (Parks et al. 2015), and only the MAGs with completeness ≥ 70% and contamination ≤ 10% were subjected to downstream analysis.

**Taxonomic profiling using the *rpS3*-based approach**

Taxonomic profiling of microbial communities was performed based on the conserved ribosomal protein rpS3 through a modified pipeline described previously (Diamond et al. 2019). Briefly, all prokaryotic *rpS3* genes were first identified across contigs of each sample with hmmsearch v3.2.1 (Eddy 2011) using a custom HMM database (Diamond et al. 2019) and then clustered at 99% similarity with USEARCH v11.0.667 (Edgar 2010), with each cluster of *rpS3* genes assigned to be a species. Among the contigs containing marker genes belonging to a *rpS3* cluster, the one that was assigned to a MAG with the largest completeness was selected as the representative sequence for each species. When all contigs corresponding to the *rpS3* cluster were not retrieved in any MAGs, the longest one was selected. To obtain the abundance of each species across samples, the clean reads were aligned against its representative sequence using Bowtie2 v2.3.5.1 (Langmead and Salzberg 2012), and then the mapped reads with ≥ 99% sequence identity were filtered and counted using the ‘depth’ module of Samtools v1.15.1 (Li et al. 2009). The final abundance of each species in a sample was calculated as the total mapped bases on the corresponding representative sequence divided by the length of the representative sequence and the total number of sequencing bases in the sample.

The taxonomy for each species was first determined by aligning the amino acids of the *rpS3* gene against a custom *rpS3* reference database using BlastP with e-value ≤ 1e-3 and percent identity ≥ 50, where each reference *rpS3* gene was derived from the RefSeq prokaryotic genome database (downloading date: 2019-07, ~27000 genomes). The taxonomy was further validated and corrected according to the rpS3 gene tree, which were built below. The amino acid sequences of all representative *rpS3* genes were aligned using MAFFT v7.427 (Katoh et al. 2005), and trimmed using trimAl v1.4.1 with the ‘-automated1’ option (Capella-Gutiérrez et al. 2009). An approximately-maximum-likelihood phylogenomic tree was built by FastTree v2.1.11 (Price et al. 2010), which is an efficient tool for inferring phylogenies with up to hundreds of thousands of species. When taxonomy assignment for species from Blastp disagrees with the phylogenetic assignment, the latter result was determined for the species. The taxonomies of the species that had no hits in the reference database and branched deeply in microbial lineages in the *rpS3* tree were designated the ‘Unassigned’ group.

The representative genomes for a subset of species were identified through the *rpS3*-containing contig sequence shared between the reconstructed MAGs and the species. Not all genomes of species were recovered using metagenome binning methods partly due to the great diversity and insufficient sequencing coverage for the samples. However, the set of representative genomes could well represent the microbial community according to two pieces of evidence: (i) the close relatedness of species evenness, species richness and Shannon diversity (Fig. S2a-c) and (ii) the high correlation of microbial composition using the Mantel test between the MAG-and species-based community (Fig. S2d). These representative genomes accounted for 24% of total species by identity, and 36% by relative abundance (Fig. S5)

**Three validation genomic dataset**

(i) The first one is a set of 345 redundant MAGs reconstructed in this study. These MAGs were independent from the 647 representative MAGs above, and were not included in any of data analyses described above. (ii) The second dataset consists of 530, 464 and 606 MAGs from freshwater, marine and alkaline habitats, respectively. These MAGs with genome completeness >= 90% and contamination <= 5% were retrieved from the Genomes from Earth’s Microbiomes (GEM) catalog for freshwater and marine environments (Nayfach et al. 2021), and from soda lakes located in regions of Russia (Vavourakis et al. 2016), China (Zhao et al. 2020) and Canada (Zorz et al. 2019) for alkaline environments. (iii) The third dataset contains 830 high-quality MAGs from freshwater environments and soda lakes with pH values available. A subset of species-level MAGs (n = 668) was retrieved from the StratFreshDB database, which is reconstructed metagenomes from freshwater lakes and ponds with pH ranging from 4.2 to 9.2 (Buck et al. 2021). The other species-level MAGs (n = 162) were reconstructed using the custom pipeline above from metagenomic samples of soda lakes with pH from 9.7 to 10.8 (Zhao et al. 2020).

Together, there were two types of genome datasets explored in the study. The first type comprised the ones reconstructed from 39 African lake samples, including the representative species-level genomes (n = 647) and the redundant genomes (345). The second type contained the ones retrieved or reconstructed from public literatures, including the genomes from freshwater or soda lakes (Zhao et al. 2020, Buck et al. 2021, Nayfach et al. 2021) and the metagenomes (Alneberg et al. 2018, Sorensen et al. 2019) . The representative genome dataset was used to establish the iEA for pH, whereas the remaining datasets were used to demonstrate the utility of iEA.

**The selection of the permutation strategy**

There are a wide range of statistical methods established to analyze the trait-environment relationships (Kleyer et al. 2012, Zelený 2018, Lepš and de Bello 2023). The community-weighted mean (CWM) method is one of the popular methods to quantify the traits at the community level, and the standard correlation test for CWM-environmental relationships has been criticized with inflated Type I errors (Peres-Neto et al. 2017). To correct for the potential Type I error rates, various statistical methods are developed subsequently, including the fourth-corner analysis with the max test, CWM-RDA (redundancy analysis), double canonical correspondence analysis, CWM/SNC (species niche centroids) approach, and the CWM method with the permutation max test (Zelený 2018). The CWM method with the permutation strategy is has been widely applied in previous studies for functional traits of plant communities along environment gradients (Zelený 2018, Miller et al. 2019, ter Braak 2019, Bricca et al. 2023).

By applying the row-and-column permutation test, we found that there were 952 KOs (14.4%) significantly associated with pH, while the remaining 5,648 KOs (85.6%) showed nonsignificant correlations. Out of the 952 KO-represented traits, we identified at least 97 KOs being well-known for their involvement in microbial pH adaptation (Krulwich et al. 2011, Banciu and Muntyan 2015). For instance, there were 22 traits associated with cytoplasmic pH homeostasis, and 21 traits relevant to the biosynthesis and uptake of compatible solutes (FigShare Table S5).

The permutation method is shown to be robust in selecting the traits with ecological meanings. We tested the robustness of the permutation method in selecting the core traits with ecological meanings stated above by reducing total number of traits included in the very beginning. Specifically, we selected four groups of KOs based on their occurrences across genomes, that is, 6,600 (occurring in all 647 species), 3,027 (occurring in > 5% species), 2,193 (10%) and 1,392 (20%) KOs. For each of four KO groups, the permutation method was applied to test their relationships with pH. We found that the set of 256 significant KOs occurring in 20% of species were consistently identified in the other three tests of different KO groups. It is the same case for the set of 411 significant KOs occurring in 10% of species identified in the other two tests of lower occurrence rates. The results clearly indicate that the permutation method is stable and robust for the variation in trait numbers.

Compared to the standard Spearman correlation test, the CWM method with the permutation strategy could better refine the KOs. With the permutation strategy used in this study, we found that there were 952 significantly pH-associated KOs. Based on the Spearman correction test, however, we found there were more KOs, that is, 1,577 KOs, showing significant relationships with lake pH (*P* < 0.05). Unexpectedly, we noted that there were 17 among the 97 KOs with well-known functions missing in the standard Spearman correlation test.

There were also a few trait-environment studies utilizing the permutation strategy along with the multiple testing correction (Dray et al. 2014). For instance, Dray et al. used the permutation strategy to analyze the trait-environment relationship, with 49,999 permutations for the 96 possible association tests (eight plant traits and 12 environment variables), and multiple testing correction using Benjamini and Hochberg (BH) method to adjust *P* values (Dray et al. 2014). This approach however will require a large number of multiple comparisons in case there are thousands of traits, and thus substantial computational resources, which could be a limitation for many researchers. The multiple testing correction usually require a higher number of permutations to detect the items with the adjusted *P* values lower than the selected significance threshold, i.e. *P* < 0.05. For example, 49,999 permutations are recommended to detect significant associations for less than 100 tests for multiple test correction in the study above(Dray et al. 2014). Considering more than 6,000 tests across two habitats in our case, at least 12,000,000 permutations are required to obtain significant items. We estimated at least three months would be needed to perform 12,000,000 permutations with a computational resource of 112 cores and 512 Gb RAM. Therefore, the permutation method combined with multiple testing correction appear challenging for very large number of KO-represented traits in microbial ecology as in our study.

Unfortunately, we found that the permutation strategy with multiple testing correction could discard nearly half of the KOs with ecological meanings in environmental adaptation. We tested the relationships between pH and the 97 KOs stated earlier (FigShare Table S5), and selected 49,999 permutations considering the *P* value adjustment with BH method. Among the 97 KOs with well-known biological functions, we however found there were only a half of these KOs kept. In the other KOs being nonsignificant and discarded, there were very important KOs, such as the one encoding the Na^+^-transporting NADH:ubiquinone oxidoreductase and the osmoprotectant transport system, which are well demonstrated to mediate microbial alkaline adaptation (Krulwich et al. 2011, Banciu and Muntyan 2015).

We further evaluated how the removal of low-TER-strength KOs (that is, the potentially bad KOs) would affect the adaptation index in case there were potential false positives in our results. Among the 952 KOs identified by the permutation strategy, the TER strength ranged from 0.524 to 0.906. We sorted the 952 KOs by the order of the TER strength, removed certain proportion (from 5% to 40%) of KOs from the lowest strength, and calculated the correlation between the new index and the original one based on all significant KOs. We found that the species adaptation index was resistant with the exclusion of these low-TER-strength KOs as shown in their correlations along the proportion of deleted KOs (Fig. S5). For instance, the Spearman correlations between species adaptation index were still larger than 0.97 for both habitats even we removed the low-TER-strength KOs up to 40% of the KOs. Therefore, the potential false positive in the current results would have no effects on the species adaption index, and thus our main findings in this study.

By reevaluating the overall results of the row-and-column permutation strategy and FDR, we would like to focus on the ecological meanings of the KOs, rather than rely solely on statistical analyses.

Noted that the non-random distribution of microbial genes along genomes may affect the identification of the relationship between the gene-represented traits and environment variables, similar as the genotype-phenotype association in the genome-wide association studies (Uffelmann et al. 2021). However, the (orthologous) gene coordinates along the genomes of phylogenetically distinct microbes are not as easily captured as in our study, due to genome evolution involving extensive gene gain and loss, pervasive recombination and horizontal gene transfer (Arnold et al. 2022). The effect should be considered in the future studies focusing on the trait-environment relationships at the species or strain levels.

Nayfach, S., S. Roux, R. Seshadri, D. Udwary, N. Varghese, F. Schulz, D. Wu, D. Paez-Espino, I. M. Chen, M. Huntemann, K. Palaniappan, J. Ladau, S. Mukherjee, T. B. K. Reddy, T. Nielsen, E. Kirton, J. P. Faria, J. N. Edirisinghe, C. S. Henry, S. P. Jungbluth, D. Chivian, P. Dehal, E. M. Wood-Charlson, A. P. Arkin, S. G. Tringe, A. Visel, H. Abreu, S. G. Acinas, E. Allen, M. A. Allen, L. V. Alteio, G. Andersen, A. M. Anesio, G. Attwood, V. Avila-Magaña, Y. Badis, J. Bailey, B. Baker, P. Baldrian, H. A. Barton, D. A. C. Beck, E. D. Becraft, H. R. Beller, J. M. Beman, R. Bernier-Latmani, T. D. Berry, A. Bertagnolli, S. Bertilsson, J. M. Bhatnagar, J. T. Bird, J. L. Blanchard, S. E. Blumer-Schuette, B. Bohannan, M. A. Borton, A. Brady, S. H. Brawley, J. Brodie, S. Brown, J. R. Brum, A. Brune, D. A. Bryant, A. Buchan, D. H. Buckley, J. Buongiorno, H. Cadillo-Quiroz, S. M. Caffrey, A. N. Campbell, B. Campbell, S. Carr, J. Carroll, S. C. Cary, A. M. Cates, R. A. Cattolico, R. Cavicchioli, L. Chistoserdova, M. L. Coleman, P. Constant, J. M. Conway, W. P. Mac Cormack, S. Crowe, B. Crump, C. Currie, R. Daly, K. M. DeAngelis, V. Denef, S. E. Denman, A. Desta, H. Dionisi, J. Dodsworth, N. Dombrowski, T. Donohue, M. Dopson, T. Driscoll, P. Dunfield, C. L. Dupont, K. A. Dynarski, V. Edgcomb, E. A. Edwards, M. S. Elshahed, I. Figueroa, B. Flood, N. Fortney, C. S. Fortunato, C. Francis, C. M. M. Gachon, S. L. Garcia, M. C. Gazitua, T. Gentry, L. Gerwick, J. Gharechahi, P. Girguis, J. Gladden, M. Gradoville, S. E. Grasby, K. Gravuer, C. L. Grettenberger, R. J. Gruninger, J. Guo, M. Y. Habteselassie, S. J. Hallam, R. Hatzenpichler, B. Hausmann, T. C. Hazen, B. Hedlund, C. Henny, L. Herfort, M. Hernandez, O. S. Hershey, M. Hess, E. B. Hollister, L. A. Hug, D. Hunt, J. Jansson, J. Jarett, V. V. Kadnikov, C. Kelly, R. Kelly, W. Kelly, C. A. Kerfeld, J. Kimbrel, J. L. Klassen, K. T. Konstantinidis, L. L. Lee, W.-J. Li, A. J. Loder, A. Loy, M. Lozada, B. MacGregor, C. Magnabosco, A. Maria da Silva, R. M. McKay, K. McMahon, C. S. McSweeney, M. Medina, L. Meredith, J. Mizzi, T. Mock, L. Momper, M. A. Moran, C. Morgan-Lang, D. Moser, G. Muyzer, D. Myrold, M. Nash, C. L. Nesbø, A. P. Neumann, R. B. Neumann, D. Noguera, T. Northen, J. Norton, B. Nowinski, K. Nüsslein, M. A. O’Malley, R. S. Oliveira, V. Maia de Oliveira, T. Onstott, J. Osvatic, Y. Ouyang, M. Pachiadaki, J. Parnell, L. P. Partida-Martinez, K. G. Peay, D. Pelletier, X. Peng, M. Pester, J. Pett-Ridge, S. Peura, P. Pjevac, A. M. Plominsky, A. Poehlein, P. B. Pope, N. Ravin, M. C. Redmond, R. Reiss, V. Rich, C. Rinke, J. L. M. Rodrigues, W. Rodriguez-Reillo, K. Rossmassler, J. Sackett, G. H. Salekdeh, S. Saleska, M. Scarborough, D. Schachtman, C. W. Schadt, M. Schrenk, A. Sczyrba, A. Sengupta, J. C. Setubal, A. Shade, C. Sharp, D. H. Sherman, O. V. Shubenkova, I. N. Sierra-Garcia, R. Simister, H. Simon, S. Sjöling, J. Slonczewski, R. S. Correa de Souza, J. R. Spear, J. C. Stegen, R. Stepanauskas, F. Stewart, G. Suen, M. Sullivan, D. Sumner, B. K. Swan, W. Swingley, J. Tarn, G. T. Taylor, H. Teeling, M. Tekere, A. Teske, T. Thomas, C. Thrash, J. Tiedje, C. S. Ting, B. Tully, G. Tyson, O. Ulloa, D. L. Valentine, M. W. Van Goethem, J. VanderGheynst, T. J. Verbeke, J. Vollmers, A. Vuillemin, N. B. Waldo, D. A. Walsh, B. C. Weimer, T. Whitman, P. van der Wielen, M. Wilkins, T. J. Williams, B. Woodcroft, J. Woolet, K. Wrighton, J. Ye, E. B. Young, N. H. Youssef, F. B. Yu, T. I. Zemskaya, R. Ziels, T. Woyke, N. J. Mouncey, N. N. Ivanova, N. C. Kyrpides, E. A. Eloe-Fadrosh, and I. M. D. Consortium. 2021. A genomic catalog of Earth’s microbiomes. Nature Biotechnology 39:499-509.

Parks, D. H., M. Imelfort, C. T. Skennerton, P. Hugenholtz, and G. W. Tyson. 2015. CheckM: assessing the quality of microbial genomes recovered from isolates, single cells, and metagenomes. Genome Research 25:1043-1055.

Peres-Neto, P. R., S. Dray, and C. J. F. ter Braak. 2017. Linking trait variation to the environment: critical issues with community-weighted mean correlation resolved by the fourth-corner approach. Ecography 40:806-816.

Price, M. N., P. S. Dehal, and A. P. Arkin. 2010. FastTree 2 – approximately maximum-likelihood trees for large alignments. PloS One 5:e9490.

Ren, M., and J. Wang. 2022. Phylogenetic divergence and adaptation of Nitrososphaeria across lake depths and freshwater ecosystems. The ISME Journal 16:1491-1501.

Sieber, C. M. K., A. J. Probst, A. Sharrar, B. C. Thomas, M. Hess, S. G. Tringe, and J. F. Banfield. 2018. Recovery of genomes from metagenomes via a dereplication, aggregation and scoring strategy. Nature Microbiology 3:836-843.

Sorensen, J. W., T. K. Dunivin, T. C. Tobin, and A. Shade. 2019. Ecological selection for small microbial genomes along a temperate-to-thermal soil gradient. Nature Microbiology 4:55-61.

ter Braak, C. J. F. 2019. New robust weighted averaging- and model-based methods for assessing trait–environment relationships. Methods in Ecology and Evolution 10:1962-1971.

Uffelmann, E., Q. Q. Huang, N. S. Munung, J. de Vries, Y. Okada, A. R. Martin, H. C. Martin, T. Lappalainen, and D. Posthuma. 2021. Genome-wide association studies. Nature Reviews Methods Primers 1:59.

Vavourakis, C. D., R. Ghai, F. Rodriguez-Valera, D. Y. Sorokin, S. G. Tringe, P. Hugenholtz, and G. Muyzer. 2016. Metagenomic insights into the uncultured diversity and physiology of microbes in four hypersaline soda lake brines. Frontiers in Microbiology 7.

Wu, Y. W., B. A. Simmons, and S. W. Singer. 2016. MaxBin 2.0: an automated binning algorithm to recover genomes from multiple metagenomic datasets. Bioinformatics 32:605-607.

Zelený, D. 2018. Which results of the standard test for community-weighted mean approach are too optimistic? Journal of Vegetation Science 29:953-966.

Zhao, D., S. Zhang, Q. Xue, J. Chen, J. Zhou, F. Cheng, M. Li, Y. Zhu, H. Yu, S. Hu, Y. Zheng, S. Liu, and H. Xiang. 2020. Abundant taxa and tavorable pathways in the microbiome of soda-saline lakes in Inner Mongolia. Frontiers in Microbiology 11:1740.

Zorz, J. K., C. Sharp, M. Kleiner, P. M. K. Gordon, R. T. Pon, X. Dong, and M. Strous. 2019. A shared core microbiome in soda lakes separated by large distances. Nature Communications 10:4230.

**Supplementary Figures**

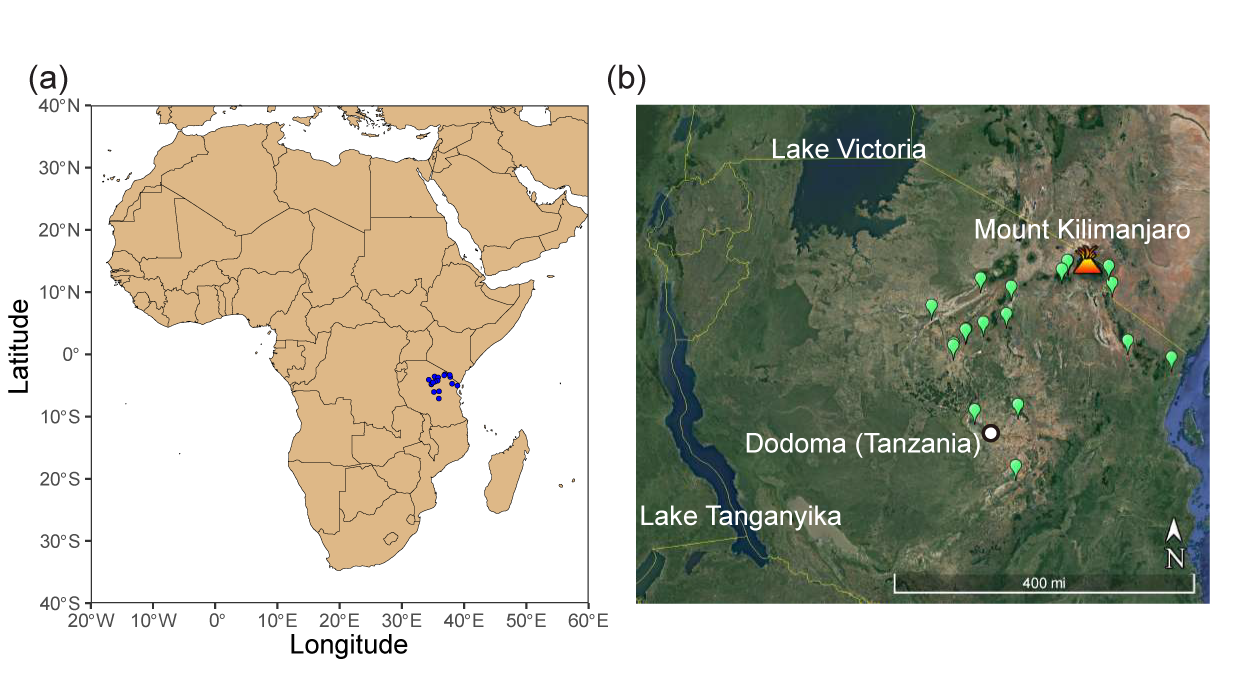

**Figure S1. The geological location of samples in lakes and reservoirs in Eastern Africa.** A total of 39 samples in the study, including 19 water and 20 sediment samples were collected from 19 lakes and reservoirs in Eastern Africa. The locations of these sites are highlighted in the map at the scale of the African continent (a) and in the view of Google Earth (b).

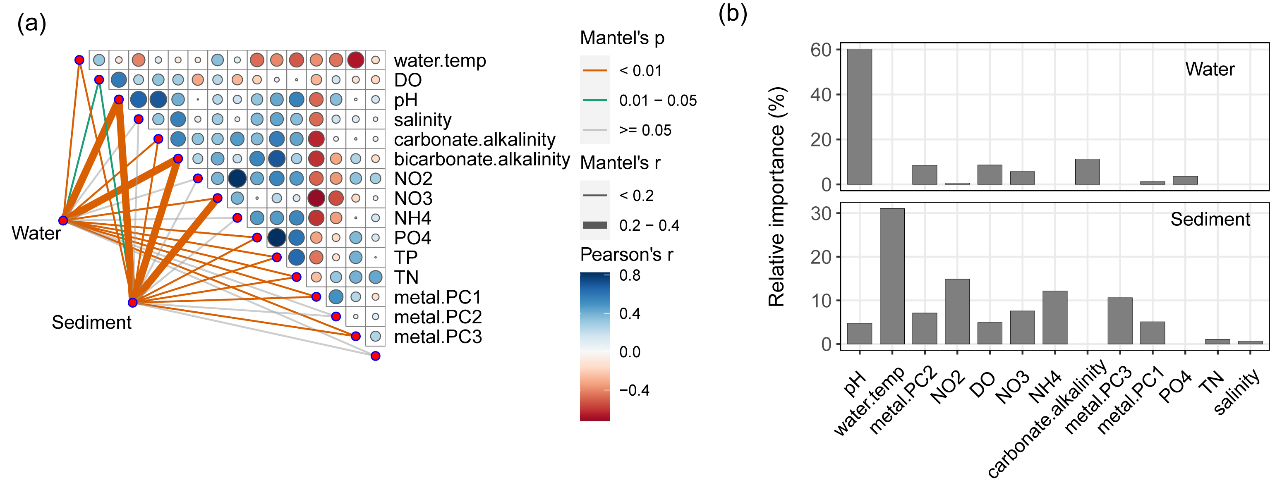

**Figure S2. The correlations between environment variables and microbial communities.** (a) Mantel test was used to analyze the correlation between a group of environmental variables and the microbial communities across water and sediments. Heatmap of the pairwise Pearson correlations between environment variables across African lakes was also shown. (b) The relative contribution of environmental factors to the variation in Shannon diversity of water and sediment communities was estimated using the random forest model.

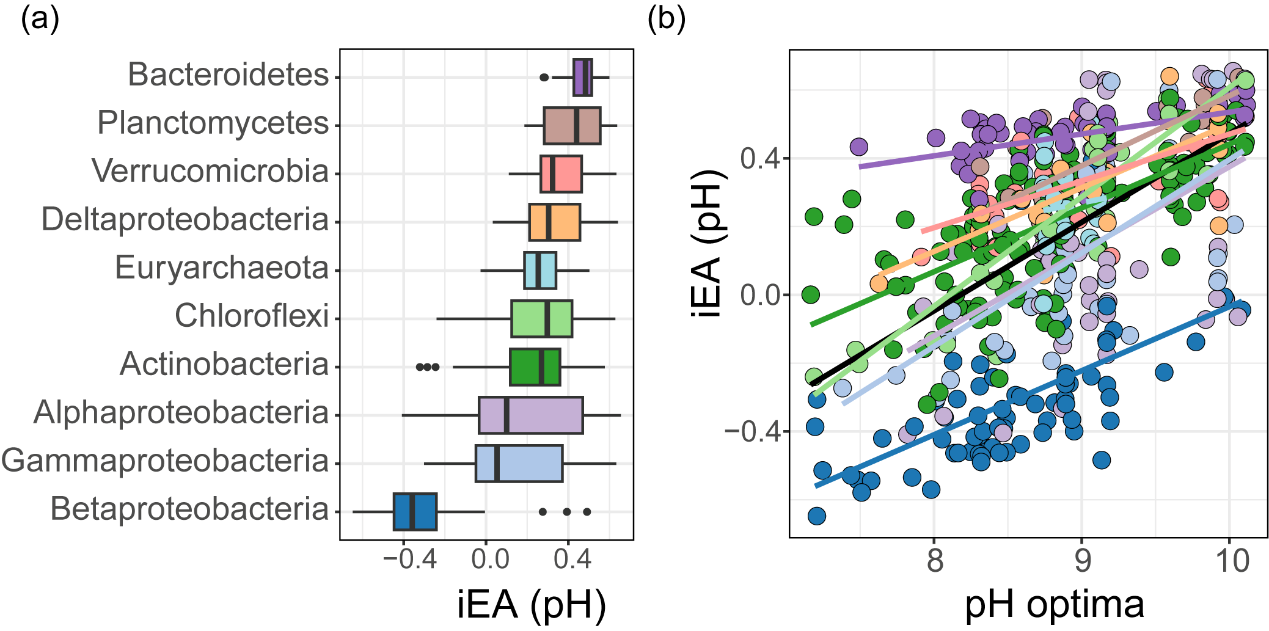

**Figure S3. The distribution of the index of environmental adaptation across phyla.** The species iEA was compared across major prokaryotic phyla (a). The relationships between iEA and species pH optima were shown at the phylum level (b). The relationship was fitted using a linear regression model with the solid line representing the significant one. Each phylum was highlighted with a distinct color. The complete statistical report for each phylum was shown as follow, Chloroflexi: *R^2^_adj_* = 0.783, *F_1,31_* = 116.38, *p* = 5.08e-12; Planctomycetes: *R^2^_adj_* = 0.623, *F_1,20_* = 35.75, *p* = 7.58e-6; Actinobacteria: *R^2^_adj_* = 0.439, *F_1,145_* = 116.19, *p* = 2.70e-20; Deltaproteobacteria: *R^2^_adj_* = 0.435, *F_1,21_* = 17.95, *p* = 3.68e-4; Gammaproteobacteria: *R^2^_adj_* = 0.411, *F_1,42_* = 31.98, *p* = 1.67e-6; Verrucomicrobia: *R^2^_adj_* = 0.382, *F_1,23_* = 15.85, *p* = 5.89e-4; Betaproteobacteria: *R^2^_adj_* = 0.331, *F_1,69_* = 35.63, *p* = 9.23e-8; Bacteroidetes: *R^2^_adj_* = 0.304, *F_1,60_* = 27.62, *p* = 2.06e-6; Alphaproteobacteria: *R^2^_adj_* = 0.191, *F_1,40_* = 10.67, *p* = 2.24e-3; Euryarchaeota: *R^2^_adj_* = -0.0406, *F_1,19_* = 0.220, *p* = 0.644.

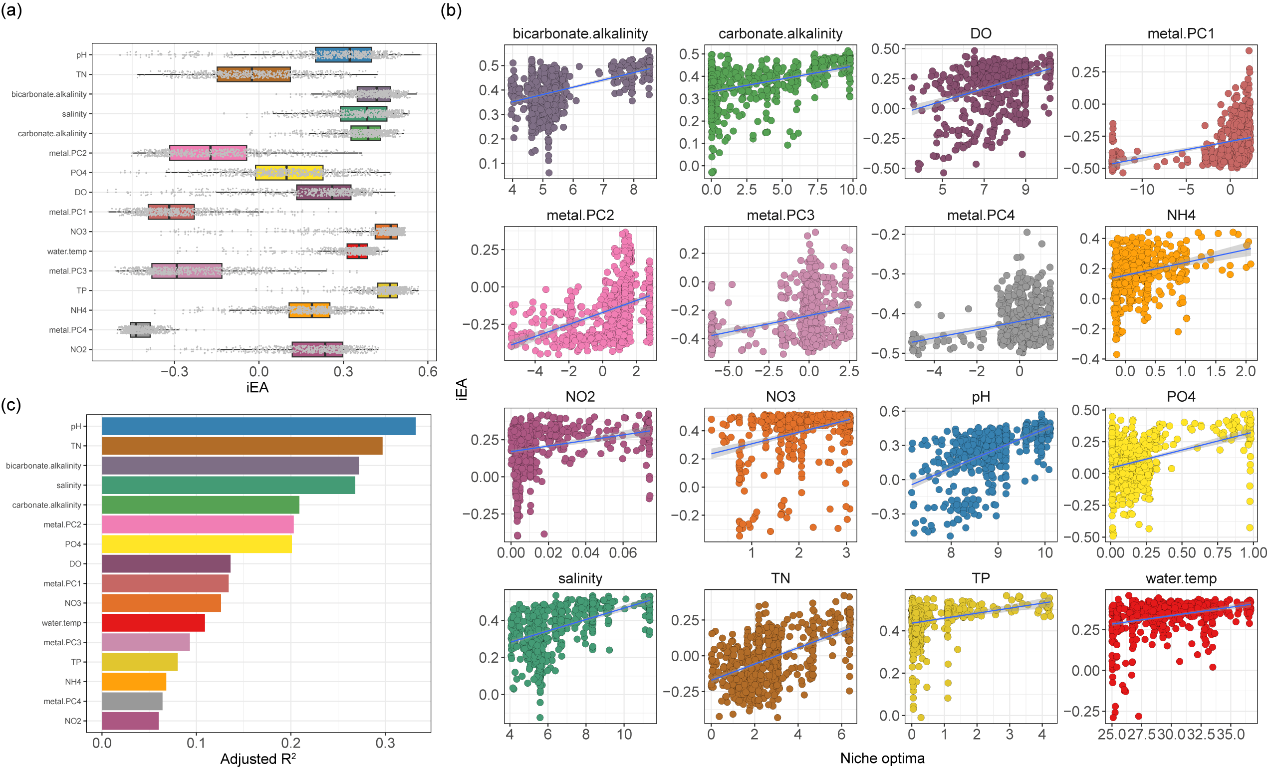

**Figure S4. The distribution and comparison of the index of environmental adaptation for all environmental factors.** The species iEA concerning all environmental factors were calculated using the same way as the iEA of pH in the study. The distribution of iEA for all environmental factors was shown in (a). Similarly, the relationships between iEA for all factors and their corresponding niche optima were also fitted using linear regression models (b). The variation in niche optima explained by the iEA was compared across all environmental factors (c).

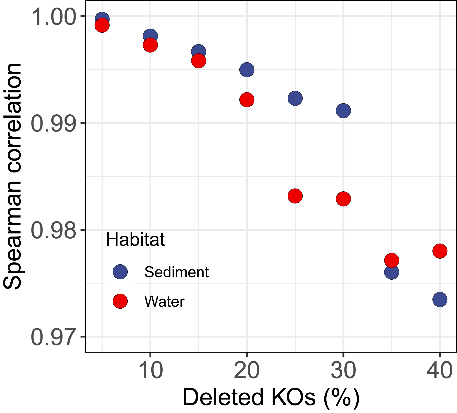

**Figure S5. The Spearman correlation between the species adaptation index along the proportion of KOs deleted from the KO pool.** We evaluated how the removal of the low-TER-strength KOs would affect the adaptation index in case there were potential false positives in the results. We sorted the 952 KOs by the order of the TER strength, removed certain proportion (from 5% to 40%) of KOs from the lowest strength, and calculated the correlation between the new index and the original one based on all significant KOs.
